# Phage protease enzymes activate CBASS antiphage immunity

**DOI:** 10.64898/2026.03.04.709575

**Authors:** Samuel J. Hobbs, Philip J. Kranzusch

## Abstract

Cyclic oligonucleotide-based antiphage signaling systems (CBASS) are immunity pathways in bacteria that use a cGAS/DncV-like nucleotidyltransferase (CD-NTase) enzyme to sense phage infection and initiate antiviral defense. Bacteria encode thousands of diverse CBASS operons, demonstrating a critical role for CD-NTase activation in controlling the prokaryotic response to viral infection. Here we discover proteolytic cleavage by phage prohead proteases as a mechanism of CD-NTase activation and demonstrate that diverse CBASS operons function as molecular sensors of protease activity. We reconstitute CBASS recognition of phage T4 infection *in vitro* and identify proteolytic cleavage of a surface exposed CD-NTase activation loop as a trigger of enzyme catalysis and nucleotide immune signal synthesis. Phage prohead proteases are sufficient to activate CBASS *in vivo* and explain how immune signaling is initiated during late stages of viral infection. Combining biochemical and structure-based phylogenetic analyses, we map activation loops in Clade A, D, and G CD-NTase enzymes and define critical residues that control CBASS recognition of distinct phage families. Our results define CBASS recognition of phage protease activity as a widespread mechanism of antiviral defense.

## Introduction

Animal, plant, and bacterial cells each synthesize specialized nucleotide-based signals to respond to viral infection and initiate antiviral defense(*1–4*). In animals, cGAS-like receptors sense infection through recognition of viral DNA or RNA and synthesize a cyclic dinucleotide signal to activate the receptor STING (stimulator of interferon genes) (*5–7*). Bacteria encode an evolutionarily related pathway, named CBASS (cyclic oligonucleotide-based antiphage signaling systems) where ancestrally related cGAS-DncV-like nucleotidyltransferase (CD-NTase) enzymes sense phage infection and synthesize cyclic di- and trinucleotide signals to activate downstream effector proteins and antiviral defense (*8–10*). Activation of CBASS immunity causes an abortive infection response, whereby bacteria become dormant or commit to cell death to restrict phage replication and prevent viral spread to neighboring cells (*8, 11, 12*). Due to this drastic fitness cost, CBASS is activated only late during the phage replication cycle and functions as a last line of antiviral defense. Consistent with this role in antiviral immunity, recent studies have identified CBASS-resistant escape mutations in phage genes including the major capsid, scaffolding proteins, and late viral non-coding RNAs associated with the final steps of virion formation (*13–15*), but the specific molecular factors recognized by most CD-NTase enzymes are unknown.

## Results

### A phage factor expressed late during infection activates CBASS signaling

To define the factors sensed by CBASS anti-phage defense we sought to biochemically reconstitute the key step of virally induced CD-NTase activation *in vitro* using the model virus *E. coli* phage T4. Phage T4 and other *Tevenvirinae* family phages encode a set of anti-CBASS (Acb) proteins dedicated to disrupting the host nucleotide signals that control CBASS anti-phage defense (*13, 16, 17*). We confirmed that an engineered phage T4 variant lacking *acb1* and *acb2* (T4 Δacb1/Δacb2) is exquisitely sensitive to an *E. coli* type II CBASS operon (*18*), demonstrating that Acb proteins are required to mask the effect of an unknown factor from phage T4 that activates CBASS (Fig. 1A,B). Inactivating mutations to the *E. coli* CD-NTase CdnG (*Ec*CdnG), regulatory Cap2/Cap3, and DNase effector Cap5 proteins each restored phage replication, supporting that phage T4 Δacb1/Δacb2 restriction occurs through canonical CD-NTase activation and downstream CBASS signaling (Fig. 1C; Fig. S1A) (*11, 17, 19, 20*). To biochemically reconstitute CD-NTase activation, we next prepared a soluble lysate from cells infected with phage T4 Δacb1/Δacb2 and mixed this sample with a separate lysate prepared from cells expressing an effector-dead variant of CBASS (Cap5 H56A). We analyzed CD-NTase activation in the resulting reconstitution reactions by developing an *in vitro* Cap5 biosensor assay that induces cleavage of an indicator dsDNA plasmid in response to the *Ec*CdnG nucleotide signal 3′2′-cGAMP (Fig. 1D; Fig. S1B,C) (*21, 22*). Using this reconstituted system, we observed that CD-NTase-dependent 3′2′-cGAMP synthesis occurs only in the presence of lysates from cells infected with phage T4 Δacb1/Δacb2 and not from uninfected cells (Fig. 1E). We tested a time-course of phage T4 Δacb1/Δacb2 infection and observed CD-NTase activation in all lysates prepared 14 minutes post infection or later (Fig. 1F). These results demonstrate that CBASS activation specifically occurs in response to a soluble factor produced during late stages of phage T4 infection.

**Figure 1:**
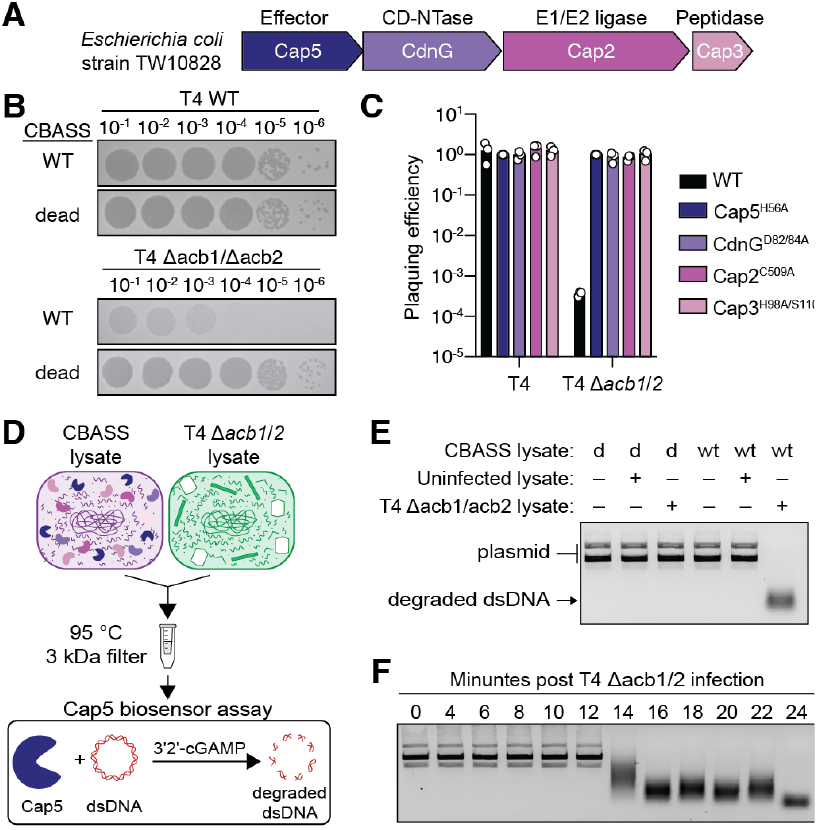
A phage factor expressed late during infection activates CBASS signaling. **(A)** Schematic of the type II CBASS operon from *E. coli* strain TW10828 used in this study. **(B)** Representative plaque assay depicting serial dilutions of WT phage T4 or phage T4 Δacb1/Δacb2 plated on cells expressing either WT or dead *Ec*CdnG D82/84A CBASS operons. **(C)** Quantification of the data in B and Figure S1A. **(D)** Experimental schematic depicting biochemical reconstitution of CBASS sensing T4 Δacb1/Δacb2 phage infection by mixing CBASS and phage-infected lysates. Activation of CBASS signaling in the reconstitution reactions is then analyzed using the Cap5-based 3′2′-cGAMP biosensor assay. **(E)** Agarose gel depicting production of 3′2′-cGAMP and cleavage of indicator dsDNA by Cap5 when CBASS lysates are mixed with T4 Δacb1/Δacb2 infected lysates. For CBASS lysates, ‘d’ indicates cells expressing operon encoding an inactive *Ec*CdnG (D82/84A) and ‘wt’ indicates cells expressing an operon encoding WT *Ec*CdnG and an inactive Cap5 (H56A). ‘–’ indicates buffer was added to the reaction and ‘+’ indicates that lysate was added to the reaction. **(F)** Agarose gel depicting production of 3′2′-cGAMP and cleavage of indicator dsDNA when CBASS Cap5 H56A lysates are mixed with T4 Δacb1/Δacb2 phage lysates harvested 14 minutes post infection or later.

### CBASS signaling is activated by proteolytic activity

Phage T4 infection in *E. coli* occurs through a well-defined sequence of molecular events. The timing of CBASS immune activation observed in our reconstitution experiments occurs immediately after the major event of phage T4 virion prohead formation at approximately 12 min post-infection (Fig. 1F) (*23*). We therefore screened candidate phage T4 proteins that are produced during the onset of virion prohead formation (*24*) and observed that expression of the viral prohead protease protein gp21 was sufficient to trigger CBASS immune activation *in vivo* (Fig. 2A; Fig. S2A,B). Surprisingly, activation of CBASS-dependent cell toxicity was dependent on the phage T4 gp21 protease active site, suggesting that *Ec*CdnG CD-NTase activation may occur in response to proteolytic activity and not simply complex formation with a specific viral factor (Fig. 2A; Fig. S2A,B). The phage T4 gp21 protease is known to be restrained by auto-proteolysis that rapidly terminates enzymatic function (*25, 26*). To test the ability of *Ec*CdnG to detect proteolysis we therefore modeled prohead protease activity *in vitro* by incubating purified *Ec*CdnG with a panel of foreign proteases of varying specificity and monitored activation using the Cap5 biosensor assay. *Ec*CdnG potently responded to foreign protease activity *in vitro* with the model proteases proteinase K, trypsin, Glu-C, and chymotrypsin each triggering production of 3′2′-cGAMP (Fig. 2B; Fig. S2C,D). Previous analyses have demonstrated that CD-NTase activation can be induced *in vitro* using elevated CD-NTase enzyme concentrations, high pH (>8.5), and/or manganese divalent cations (*9, 11, 21, 27–29*). Notably, we observed that protease-dependent activation of *Ec*CdnG was >50-fold more potent than previously developed artificial reaction conditions (Fig. 2C,D). Together, these results demonstrate that foreign protease activity is sufficient to activate *Ec*CdnG to catalyze nucleotide immune signal synthesis.

**Figure 2.**
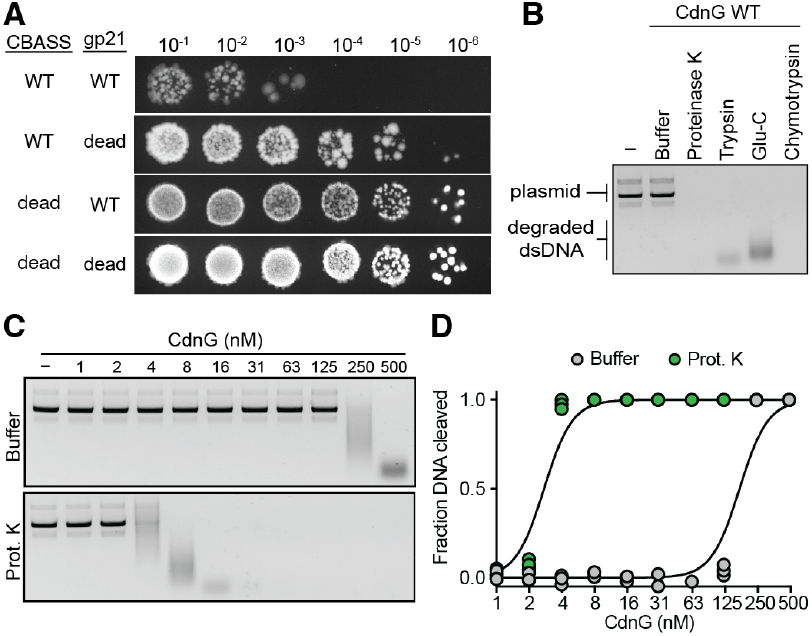
CBASS signaling is activated by proteases. **(A)** Representative toxicity assay depicting serial dilutions of *E. coli* co-transformed with the indicated CBASS operon and T4 prohead protease gp21 construct and plated on LB supplemented with 0.02% arabinose and 1 μM IPTG. For CBASS, ‘dead’ refers to *Ec*CdnG D82/84A mutations and for T4 gp21 ‘dead’ refers to H85A/S140A mutations. **(B)** Aga-rose gel depicting the Cap5 activation and cleavage of indicator dsDNA when *Ec*CdnG is treated with the indicated protease. **(C)** Representative agarose gel depicting activation of Cap5 and cleavage of indicator dsDNA when titrating amounts of *Ec*CdnG are treated with proteinase K. **(D)** Quantification of the data in C.

### Cleavage of an unstructured loop activates *Ec*CdnG

To define how foreign proteases induce CD-NTase activation we next monitored *Ec*CdnG protein stability during activation of 3′2′-cGAMP synthesis. In the presence of trypsin, activation of *Ec*CdnG 3′2′-cGAMP synthesis co-occurred with cleavage of *Ec*CdnG into an ~28 kDa protein species (Fig. 3A–C). We compared the AlphaFold3-predicted structure of *Ec*CdnG to experimentally determined structures of CD-NTase enzymes (9, 27, 28, 30–32) and identified an ~30 amino acid surface-exposed, disordered loop in the center of the canonical CD-NTase bi-lobed enzyme architecture where cleavage would result in production of an ~28 kDa product (Fig. 3A). Substitution of all four Arg/Lys trypsin cleavage sites to glutamine within this disordered *Ec*CdnG loop (K184Q, K197Q, K214Q, K218Q) abolished the ability of *Ec*CdnG to produce nucleotide immune signals in response to trypsin protease activity (Fig. 3A–C). Notably, the trypsin-blind mutant *Ec*CdnG enzyme retained the ability to sense proteinase K protease activity, demonstrating that the mutated *Ec*CdnG enzyme is functional and could still respond to proteases with alternative cleavage-site specificities.

**Figure 3.**
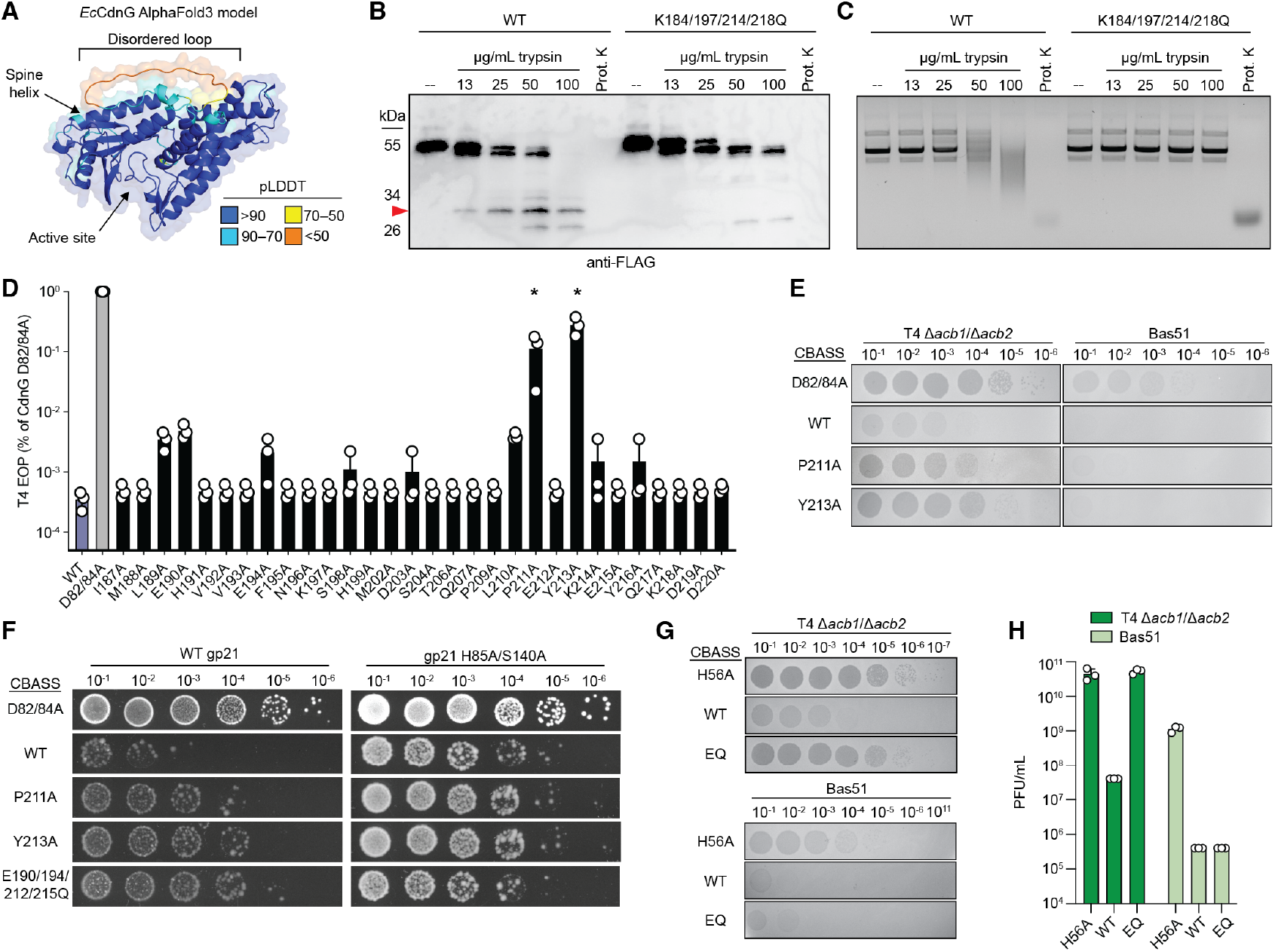
Cleavage of an unstructured loop activates *Ec*CdnG. **(A)** AlphaFold3 model of *Ec*CdnG highlighting the unstructured loop and canonical features of CD-NTase proteins. Model is colored by pLDDT confidence score. The flexible C-terminal tail of *Ec*CdnG was removed for clarity. **(B)** Representative anti-FLAG western blot of either wild-type or trypsin-blind 3×FLAG-*Ec*CdnG treated with titrating amounts of trypsin or proteinase K. Red arrow indicates ~28 kDa species that co-occurs with 3′2′-cGAMP synthesis. **(C)** Agarose gel depicting Cap5 activation and cleavage of indicator dsDNA when either wild-type or trypsin-blind 3×FLAG-*Ec*CdnG is treated with titrating amounts of trypsin or proteinase K. **(D)** Efficiency of plaquing (EOP) of T4 Δacb1/Δacb2 phage plated on cells encoding CBASS operons with the indicated *Ec*CdnG mutation. Data are normalized to the *Ec*CdnG D82/84A dead operon. **(E)** Representative plaque assays of serial dilutions of T4 Δacb1/Δacb2 phage or Bas51 phage plated on cells encoding the indicated *Ec*CdnG point mutations. **(F)** Representative toxicity assay depicting serial dilutions of *E. coli* co-transformed with CBASS operons encoding the indicated *Ec*CdnG mutations and either wild type or catalytically dead T4 prohead protease plated on 0.02% arabinose and 1 μM IPTG. **(G)** Representative plaque assays of T4 Δacb1/Δacb2 phage or Bas51 phage plated on cells encoding either inactive Cap5 (H56A), wild type, or *Ec*CdnG E190Q, E194Q, E212Q, E215Q (EQ) CBASS operons. **(H)** Quantification of the data in G.

The phage T4 gp21 protease is known to preferentially cleave target substrates C-terminal of glutamate residues within I/LxE motifs (26, 33). The *Ec*CdnG disordered loop contains an LPE motif, consistent with the ability to detect gp21 protease activity during phage T4 infection. We used alanine-scanning mutagenesis to disrupt the *Ec*CdnG disordered loop in vivo and observed that mutations to P211 or the adjacent residue Y213 in the LPE motif attenuated >100-fold the ability of CBASS to defend against phage T4 Δacb1/Δacb2 infection (Fig. 3D,E; Fig. S3A). To further define the ability of this disordered loop to control the specificity of *Ec*CdnG phage recognition, we screened the BASEL phage collection for additional susceptible phages and identified that the *Ec*CdnG CBASS operon successfully defends against phages Bas48–59 of the Vequintavirinae family (Fig. S3B) (34). *Ec*CdnG mutations P211A and Y213A disrupted CBASS defense against phage T4 but not against phage Bas51, demonstrating that these mutations alter the specificity of CBASS phage recognition and do not compromise overall immune function (Fig. 3E; Fig. S3A). Co-expression of phage T4 gp21 activated wildtype CBASS but failed to induce toxicity of *Ec*CdnG P211A or Y213A mutant operons, further confirming that mutations directly affect gp21-dependent activation of CBASS in vivo (Fig. 3F; Fig. S3C,D). Interestingly, mutation of the *Ec*CdnG residue E212 that resides within the putative LPE motif was not sufficient to abolish defense (Fig. 3D,E; Fig. S3A). However, the phage T4 gp21 protease is known to more weakly target noncanonical xxE motifs (35). We observed that an *Ec*CdnG variant with all glutamate residues in the disordered loop mutated to glutamine (E190Q, E194Q, E212Q, E215Q) no longer was able to respond to gp21 expression or defend against phage T4 replication but retained the ability to restrict phage Bas51 (Fig. 3F–H; Fig. S3C,D). Together, these data demonstrate that *Ec*CdnG contains a surface-exposed activation loop that functions as a molecular tripwire to sense phage protease function.

### CD-NTase proteins use cleavage of activation loops as a widespread mechanism of viral recognition

CD-NTases comprise a diverse family of thousands of signaling enzymes in CBASS immunity that segregate into distinct clades designated A–H (9, 36). To determine if sensing protease activity is common throughout CBASS immunity, we clustered CD-NTase protein sequences from each clade by identity and combined analysis of experimental and AlphaFold3-predicted structures to search each cluster for conserved patterns of potential activation loops. These analyses revealed that ~24% of all CBASS operons contain a CD-NTase with a large surface-exposed activation loop in the same three-dimensional location as the activation loop of *Ec*CdnG (Fig. 4A). Activation loops are present in nearly all CD-NTase sequences in Clades A and G, and ~18% of the sequences in Clade D, suggesting these CBASS operons may share a unified mechanism of immune activation (Fig. 4A). Structure-guided alignments of diverse representative CD-NTases reveal that within each clade, the activation loops contain limited sequence similarity but are framed by the same conserved secondary structure elements. To determine if diverse CD-NTases with putative activation loops also respond to foreign protease activity, we purified 12 proteins representing each major cluster in Clades A, D, and G and quantified the ability of these enzymes to signal in response to proteinase K (Fig. 4B; Fig. S4A). Remarkably, for the CD-NTase clade clusters predicted to contain activation loops, 7 of the 9 representative enzymes sensed protease function and catalyzed exceptionally robust nucleotide immune signal synthesis (Fig. 4B). CD-NTase enzymes responsive to protease function notably include the major model operons used to define the function of CBASS immunity *Vibrio cholerae* DncV and *Enterobacter cloacae* CdnD (8, 9, 11, 19, 37). In contrast, 3 of 3 CD-NTases from the Clade D clusters lacking predicted activation loops all failed to respond to protease function as expected (Fig. 4B). Together, these findings demonstrate proteolytic cleavage of CD-NTase activation loops is a widespread mechanism of activation shared across diverse bacteria and define detection of phage proteases as a major trigger controlling CBASS immune signaling.

**Figure 4.**
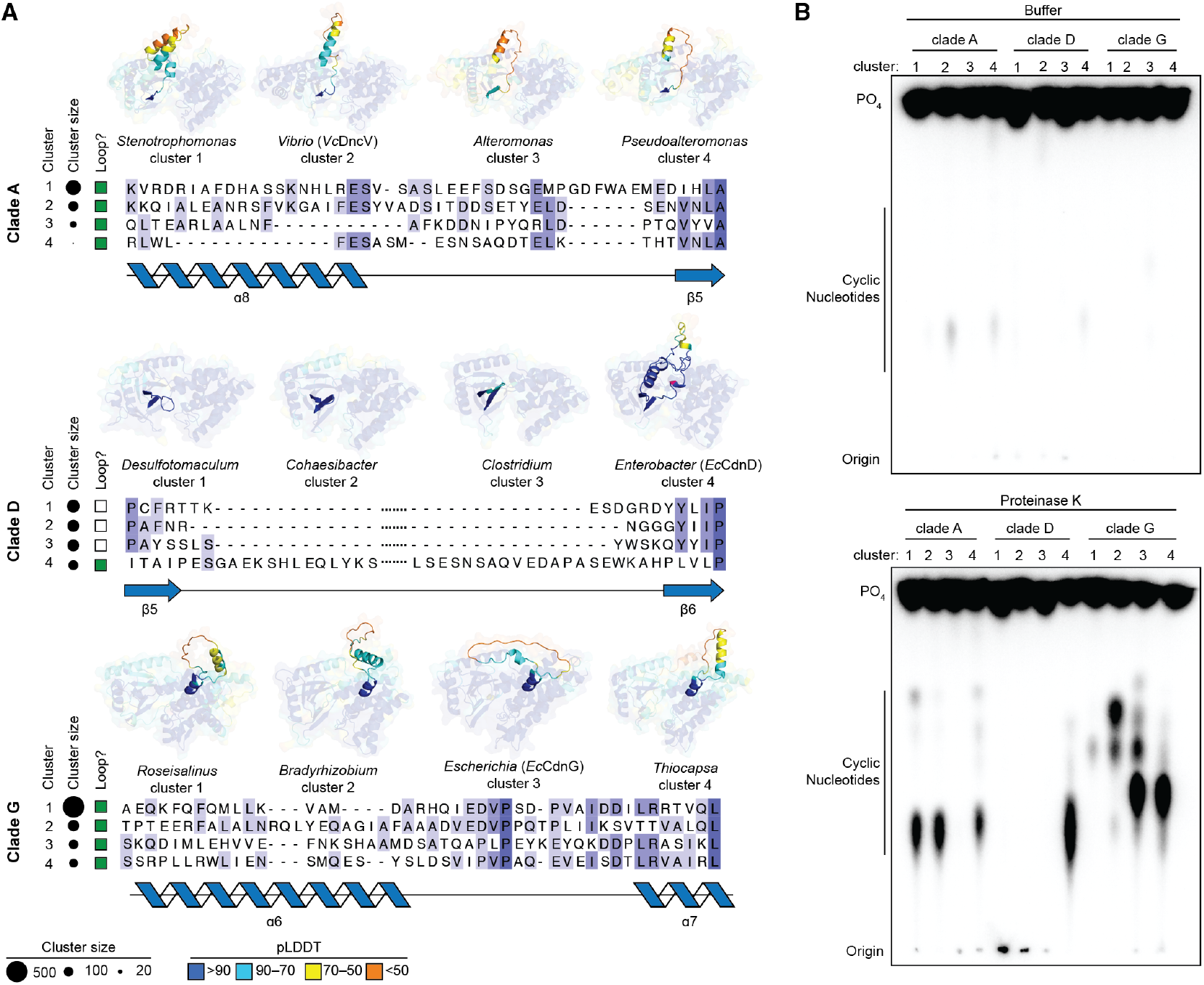
CD-NTase proteins use cleavage of activation loops as a widespread mechanism of viral recognition. **(A)** AlphaFold3 models and sequence alignments of representatives from the 4 largest clusters of CD-NTases in clade A, D, or G. The strength of shading in the sequence alignments indicates the degree of residue conservation. **(B)** Representative TLC assays depicting proteinase K-dependent activation of nucleotide immune signal synthesis by the CD-NTase cluster representatives shown in A.

## Discussion

This study identifies viral protease activity as a major trigger of CBASS anti-phage defense. The phage T4 prohead protease gp21 is sufficient to activate CBASS *in vivo*, and we show that CD-NTase enzymes sensitively respond to proteolytic cleavage as a direct biochemical trigger *in vitro*. Mechanistically, our data support a model in which a disordered CD-NTase activation loop acts as a protease-sensitive regulatory element that once cleaved enables nucleotide immune signal synthesis. We further show that activation loop architectures are conserved across >1,300 CD-NTase enzymes in multiple clades, suggesting that sensing of phage maturation through detection of protease activity is a widely conserved activating signal in CBASS immunity. CD-NTase activation loop cleavage may facilitate enzyme oligomerization or release of inhibitory molecules like folate metabolites known to restrain CBASS activation (31, 38, 39), but defining the structural transitions linking activation loop cleavage to CD-NTase catalytic function will be an important goal of future research. Protease-dependent activation aligns with the observation that CBASS acts at a late timepoint during infection and provides a mechanistic rationale to explain why phage maturation factors are enriched for CBASS escape mutations (8, 13–15). The type II CBASS components Cap2 and Cap3 are dispensable for protease-dependent activation *in vitro* but are required for phage restriction in cells, suggesting CBASS accessory proteins may function in localization or scaffolding roles to position CD-NTase enzymes to sense protease activity and virion maturation. Finally, our results parallel protease-dependent activation mechanisms in animal innate immunity including NLRP1 and CARD8 inflammasome recognition of viral protease function (40–42). Collectively, these findings reveal a major mechanism of viral sensing in CBASS immunity and define viral protease activity as a conserved foreign signal that activates anti-viral immunity in multiple kingdoms of life.

## Acknowledgements

We are grateful to members of the Kranzusch laboratory for helpful comments and discussion. We thank A. Harms for providing aliquots of the BASEL phage collection. This paper was typeset with the bioRxiv word template by @Chrelli: www.github.com/chrelli/bioRxiv-word-template

## Author contributions

The study was designed and conceived by SJH and PJK. All experiments were performed by SJH. The manuscript was written by SJH and PJK.

## Competing interest statement

The authors declare no competing interests.

## Materials and Methods

### Bacterial strains and phages

*E. coli* strain MG1655 (ATCC 47076) was used for all experiments involving phage infections, *E. coli* strain Top10 (Thermo) was used for all cloning steps, and *E. coli* strain BL21 (DE3) RIL (Agilent) was used for protein expression. The generation of phage T4 Δacb1/Δacb2 has been described previously (18). Briefly, premature stop codons were introduced into the acb1 and acb2 loci using a Cas13-based selection method as described previously(43). The BASEL phage collection was a generous gift from Dr. Alexander Harms (34).

### Cloning and plasmid construction

The native operon encoding Cap5-CdnG-Cap2-Cap3 CBASS (NCBI accession: NZ_AELC00000000.1; coordinates 2940175–2944853) was PCR amplified from existing plasmids described previously (11) and cloned into an arabinose inducible pBAD vector previously used in CBASS phage defense assays (14) using the NEBuilder HiFi Assembly Master Mix (NEB). The native operon sequence encodes a GTG start codon that was replaced with ATG to ensure proper expression in *E. coli* MG1655. Inactivating point mutations in each gene (CdnG: D82/84A; Cap5: H56A; Cap2:C509A; Cap3:H98A/S110A) were generated by site-directed mutagenesis using polymerase chain reaction (PCR) primers encoding the desired mutation and the wild type operon as a template. The resulting PCR products were then assembled into pBAD vectors using the NEBuilder HiFi Assembly Master Mix (NEB). For alanine scanning of the *Ec*CdnG activation loop, each loop residue was replaced with a GCG alanine codon and plasmids were synthesized by Twist Bioscience. The prohead protease gene of T4 (gp21; NCBI accession: NP_049785.1) was PCR amplified from the T4 genome and cloned into a custom pET-based expression vector with inducible expression controlled by Isopropyl β-D-1-thiogalactopyranoside (IPTG), a p15a origin of replication, and a chloramphenicol resistance gene. Synthetic DNA encoding a catalytic dead version of gp21 (H85A/S140A) was ordered from Integrated DNA Technologies (IDT) and cloned into the same vector. For protein expression and purification, genes encoding Cap5 or CD-NTases were synthesized (IDT) and cloned into a custom pET-based expression vector containing either an amino-terminal 6×His-SUMO2 (*Ec*Cap5, *Ec*CdnG, clade A clusters 1–4, clade D clusters 2–4, and clade G clusters 2– 4), or an amino-terminal 6×His-MBP tag (clade D cluster 1, clade G cluster 1) described previously (44).

### Phage infections and plaque assays

To amplify phage T4 Δacb1/Δacb2 and BASEL phage stocks, *E. coli* MG1655 was streaked onto a Luria Broth (LB) agar plate, incubated at 37°C overnight, and a single colony was inoculated into 3 mL LB for overnight growth at 37°C with shaking. Overnight cultures were then diluted 1:100 into 5 mL LB and grown at 37°C with shaking for approximately 1 hour until the culture reached an OD600 of approximately 0.15 and 25–50 µl of existing phage stocks (at least 106 pfu/ml) were added to the culture. Infected cultures were then incubated at 37°C with shaking for 2–6 hours until culture collapse, passed through a 0.22 µm syringe filter and stored at 4°C. To measure phage defense, *E. coli* MG1655 cells were transformed with an individual CBASS plasmid, plated on LB agar plates containing ampicillin (100 µg/ml, GoldBio) and incubated overnight at 37°C. A single colony was then picked into LB broth containing ampicillin and grown for 5–6 hours at 37°C with shaking until OD600 reached at least 1.0. LB top agar containing 0.5% (w/v) low melting point agarose (ThermoFisher) was heated to 100°C in a microwave and then placed in a 42°C water bath for 30 minutes before addition of L-arabinose (Sigma) to a final concentration of 0.02% (w/v). Five hundred µl of *E. coli* cultures were then mixed with 4 ml of melted top agar and immediately poured onto pre-warmed LB-ampicillin agar plates and allowed to solidify for 20 minutes at room temperature. Once solidified, 2.5 µl of serial dilutions of high-titer phage stocks were spotted onto the top agar and allowed to dry uncovered at room temperature for 30 minutes. Plates were then covered and incubated inverted at 30°C overnight and phage titers were determined by counting individual plaques. For active CBASS operons, titers were determined by counting a single plaque at the lowest dilution where inhibition of bacterial growth was observed.

### Recombinant protein expression and purification

*E. coli* BL21(DE3) RIL cells were transformed with expression constructs, plated on minimal glucose media (MDG) (1.5% Bacto agar, 0.5% glucose, 25 mM Na2HPO4, 25 mM KH2PO4, 50 mM NH4Cl, 5 mM Na2SO4, 0.25% aspartic acid, 2–50 µM trace metals, 100 µg/ml ampicillin, 34 µg/ml chloramphenicol) and incubated overnight at 37°C. Three colonies were then picked into 30 ml of liquid MDG media and grown overnight at 37°C with shaking. Overnight MDG cultures were then diluted 1:100 into 2 liters of M9ZB media (47.8 mM Na2HPO4, 22 mM KH2PO4, 18.7 mM NH4Cl, 85.6 mM NaCl, 1% casamino acids, 0.5% glycerol, 2 mM MgSO4, 2–50 µM trace metals) supplemented with ampicillin (100 µg/ml) and chloramphenicol (34 µg/ml), grown at 37°C with shaking for approximately 5 hours until OD600 reached 2.0, chilled in an ice bath for 20 minutes before addition of IPTG (GoldBio) to a final concentration of 0.5 mM, and incubated at 16 °C with shaking overnight. Cultures were then centrifuged at 4,000 × g and pellets were resuspended in lysis buffer (20 mM HEPES-KOH pH 7.5, 400 mM NaCl, 10% glycerol, 30 mM imidazole, 1 mM DTT). Resuspended cells were then lysed by sonication using a Q500 sonicator (QSonica) with a 1/2 inch probe at 70% amplitude using a 10 second pulse followed by 20 second rest for a total of 5 minutes of sonication. Lysates were centrifuged 30 minutes at 50,000 × g at 4°C to clear out insoluble components. Clarified lysates were flowed over 8 ml of Ni-NTA resin (Qiagen) followed by washing with 30 ml lysis buffer, 70 ml wash buffer (20 mM HEPES-KOH pH 7.5, 1 M NaCl, 10% glycerol, 30 mM imidazole, 1 mM DTT), 20 ml lysis buffer, and eluted with 20 ml elution buffer (20 mM HEPES-KOH pH 7.5, 400 mM NaCl, 10% glycerol, 300 mM imidazole, 1 mM DTT). For constructs containing a SUMO2 tag, eluted samples were then treated with recombinant human SENP2 protease to cleave the SUMO2 tag as described previously (44) and placed into dialysis tubing with a 14 kDa molecular weight cutoff and dialyzed in dialysis buffer (20 mM HEPES-KOH pH 7.5, 250 mM KCl, 10% glycerol, 1 mM DTT). Dialyzed samples were then concentrated using 10 kDa molecular weight cutoff centrifugal concentrators (Millipore Sigma) and purified further by size exclusion chromatography using a Superdex 75 column (Cytiva) using gel filtration buffer (20 mM HEPES-KOH pH 7.5, 250 mM KCl, 10% glycerol, 1 mM TCEP). Fractions containing the purified proteins were then collected and concentrated to ~500 µl and concentrations were determined by measuring A280 on a nanodrop (Denovix). Concentrated proteins were then aliquoted and flash frozen in liquid nitrogen and stored at −80°C.

### Lysate mixing reactions

To generate CBASS lysates, *E. coli* MG1655 cells were transformed with the pBAD plasmid encoding the Cap5 H56A mutant operon and plated on LB agar plates containing ampicillin (100 µg/ml). A single colony was then picked into M9ZB media (described above) supplemented with ampicillin (100 µg/ml) and grown overnight at 37°C with shaking. Overnight cultures were diluted 1:100 into 600 ml M9ZB media supplemented with ampicillin (100 µg/ml) and 0.02% L-arabinose (w/v) and grown for 1.5 hours until OD600 reached 0.15. Cells were divided into 15 ml aliquots and centrifuged at 3,200g for 10 minutes at 4°C and cell pellets were flash frozen in liquid nitrogen and stored at −80°C. To generate phage infected lysates, *E. coli* MG1655 cells were streaked onto LB agar plates and a single colony was picked into LB broth for overnight growth at 37°C with shaking. Overnight cultures were diluted 1:100 into 500 ml LB broth and grown at 37°C with shaking until OD600 reached 0.2. Cultures were then infected with phage T4 Δacb1/Δacb2 at a multiplicity of infection of 3. After 20 minutes, infected cells were divided into 5 ml aliquots, centrifuged at 3,200 × g for 10 minutes at 4°C, and cell pellets were immediately flash frozen in liquid nitrogen and stored at −80°C.

To generate lysates, CBASS pellets were resuspended in 5 ml of CBASS lysis buffer (20 mM HEPES-KOH pH 7.5, 150 mM KCl, 5 mM MgCl2, 1 mM DTT, 250 µM ATP, 250 µM GTP) and lysed by sonication using a Q500 sonicator (QSonica) with a 1/8 inch microtip at 25% amplitude for a total of 6 seconds. Phage infected pellets were resuspended in 5 ml of phage lysis buffer (20 mM HEPES-KOH pH 7.5, 150 mM KCl, 5 mM MgCl2, 1 mM DTT, 1% NP-40 alternative) and lysed by sonication using the same protocol as used for CBASS pellet lysis. Following lysis, 100 µl of CBASS lysate was then mixed with 50 µl of phage infected lysate and incubated at 30°C for 1 hour, heated to 95°C for 5 minutes, and filtered through a 3 kDa molecular weight cutoff filter to remove cellular debris. Following filtration, 2 µl of the 3 kDa flow through was used as input to the Cap5 biosensor assay (described below).

### Cap5 Biosensor Assay

Purified Cap5 was diluted to 1.11 µM in 18 µl reaction buffer (20 mM HEPES-KOH pH 7.5, 75 mM NaCl, 5 mM MgCl2, 1 mM DTT, 15 ng/µl pGEM9Z plasmid DNA (Promega)) and mixed with 2 µl of a panel of synthetic cyclic di- and tri-nucleotides (Biolog) (Fig. S1), mixed lysate reconstitution reactions (Fig. 1), or purified CdnG-protease reactions (described below) (Fig. 2, Fig. 3, Fig. S2) in a total volume of 20 µl. Cap5 reactions were then incubated at 37°C for 1 hour, heat inactivated at 95°C for 5 minutes, and stored at 4°C until analyzed by gel electrophoresis. 20 µl reactions were then mixed with 4 µl of 6× Green DNA loading dye (GoldBio) and the entire sample was loaded into wells of a 1% agarose gel containing ethidium bromide (0.5 µg/ml) and ran for 20 minutes at 150 V. Agarose gels were then visualized using a ChemiDoc MP imaging system (Bio-Rad) and cleavage of plasmid dsDNA was quantified using Fiji (45).

### Cell toxicity assay

*E. coli* MG1655 cells were transformed with pBAD plasmids encoding either wild type or *Ec*CdnG D82/84A CBASS operons together with pET plasmids encoding either wild type or catalytically inactive (H85A/S140A) T4 gp21 and plated on LB agar plates supplemented with ampicillin (100 µg/ml) and chloramphenicol (34 µg/ml). Single colonies were then picked into 3 ml LB broth supplemented with ampicillin (100 µg/ml), chloramphenicol (34 µg/ml), and 1% glucose (w/v) and grown at 37°C with shaking for 5–6 hours until OD600 reached at least 1.0. Cells were then pelleted by centrifugation at 3,200g for 10 minutes at 4°C and resuspended in 10 ml sterile phosphate buffered saline (PBS). Cells were washed a total of two times and resuspended in 3 mL PBS. 5 µl of 10-fold serial dilutions were then plated on LB agar plates supplemented with ampicillin (100 µg/ml) and chloramphenicol (34 µg/ml) and either 1% glucose to repress expression, or 0.02% L-arabinose and 1 µM IPTG to induce expression of both CBASS and gp21. Spotted dilutions were allowed to dry at room temperature for 20–30 minutes before incubation overnight at 30°C and imaging on a ChemiDoc MP imager.

### Protease treatment of CD-NTases

Lyophilized trypsin (Promega) and Glu-C (Promega) were resuspended in protease resuspension buffer (150 mM NaCl, 20 mM HEPES-KOH pH 7.5, 5 mM MgCl2, 1 mM TCEP) to a final concentration of 0.1 mg/ml. Chymotrypsin (Promega) was resuspended in 1 mM HCl to a concentration of 1 mg/ml and then diluted to 0.1 mg/ml in protease resuspension buffer. Proteinase K (NEB) was diluted to 0.4 µg/ml in protease resuspension buffer. Thermolabile proteinase K (NEB) was diluted to 0.4 µg/ml in protease resuspension buffer and either kept on ice or heat inactivated at 55 °C for 10 minutes per manufacturer’s instructions. Purified *Ec*CdnG constructs were diluted to a final concentration of 50 nM (unless otherwise indicated) in 100 µl reaction buffer (20 mM HEPES-KOH pH 7.5, 10 mM MgCl2, 50 mM KCl, 1 mM DTT, 2 mM ATP, 2 mM GTP) and mixed with 50 µl of the indicated protease. Reactions were incubated at 37°C for 1 hour, heated to 95°C for 5 minutes, spun through a 3 kDa filter and analyzed by the Cap5 biosensor assay. For testing protease-dependent activation of diverse CD-NTase clusters, serial 10-fold dilutions of purified CD-NTases were incubated in reaction buffer in the absence of any proteases and tested for activity using thin-layer chromatography (described below). The highest concentration at which CD-NTases were inactive was then used to test for proteinase K-dependent activation as described above (Table S1).

### Western blots

Purified amino-terminally 3×FLAG-tagged wild type or K184/197/214/218Q *Ec*CdnG proteins were diluted to 50 nM in 100 µl reaction buffer (20 mM HEPES-KOH pH 7.5, 10 mM MgCl2, 50 mM KCl, 1 mM DTT, 2 mM ATP, 2 mM GTP) and mixed with 50 µl of trypsin at the indicated concentration and incubated at 37°C. After 5 minutes, 0.5 µl 1M phenylmethylsulfonyl fluoride (PMSF) in dimethyl sulfoxide was added (final concentration = 3.3 mM) to inhibit trypsin activity and the reactions were incubated at 37°C for an additional 55 minutes to allow continued synthesis of 3′2′-cGAMP and then heated to 95°C for 5 minutes. Following heat inactivation, 10 µl of the reaction was mixed with 5 µl of 3× blue protein loading dye (NEB) and 10 µl of this mixture was run on a 15% SDS-PAGE gel at 200 V for 40 minutes. Proteins were then transferred to a poly(vinylidene fluoride) (Bio-Rad) membrane and non-specific binding was blocked by incubating the membrane in Tris-buffered saline with 0.1% Tween-20 (TBST) containing 5% (w/v) powdered milk (Nestle) for 30 minutes at room temperature. The membrane was then incubated in 5 ml TBST + 1% milk with 1 µl horseradish peroxidase-tagged anti-FLAG M2 antibody (Cell Signaling) for 16 hours at 4°C, followed by three washes in TBST at room temperature before bands were detected using the SuperSignal Pico Chemiluminescent substrate (Thermo) on a ChemiDocMP imager (Bio-Rad).

### Thin layer chromatography

CD-NTase reactions were performed as described above and analyzed by thin-layer chromatography as described previously (9). Briefly, reactions were prepared using the CD-NTase concentrations listed in Table S1 in reaction buffer (20 mM HEPES-KOH pH 7.5, 10 mM MgCl2, 50 mM KCl, 1 mM DTT, 2 mM ATP, 2 mM GTP, 2 mM CTP, 2 mM UTP) containing either protease resuspension buffer or proteinase K (0.4 µg/ml) supplemented with trace amounts of α-32P-labeled NTPs (PerkinElmer). Reactions were incubated at 37°C for 2 hours and then treated with Quick CIP (calf intestinal phosphatase, NEB) for 30 minutes at 37°C to remove unreacted NTPs, and 0.5 µl was spotted onto a 20 cm × 20 cm PEI-cellulose TLC plate (SigmaAldrich) and allowed to dry at room temperature for 5 minutes. Once dried, plates were placed into a running buffer of 1.5 M KH2PO4 (pH 3.8) for 1.5 hours until the solvent reached approximately 3 cm from the top of the plate. Plates were then allowed to dry at room temperature for 1.5 hours and exposed to a storage phosphor screen (Cytiva) overnight and imaged using a Typhoon Trio Variable Mode Imager System (GE Healthcare).

### CD-NTase sequence analysis

CD-NTase sequences for each clade were collected from Supplementary Table 2 from Whiteley et al, Nature 2019 (9). Sequences were then input into mmseqs2 (46) using a minimum sequence identity of 0.3 and a minimum alignment coverage of 0.8 on the MPI Bioinformatics Toolkit web server (47). A representative sequence from the 4 largest clusters from each clade was chosen at random for structure prediction, unless a previously studied CD-NTase was present within that cluster. Sequence alignments of the 4 representative sequences for each cluster was performed using the PROMALS3D web server (48) and alignment figures were generated using Jalview (49).

### AlphaFold3 structure prediction

Full length sequences of the indicated CD-NTase (Table S1) were used for structure prediction using the AlphaFold3 web server (alphafoldserver.com) (50). The highest confidence model for each cluster was used for loop analysis and figures were created using PyMOL v 3.10. Low confidence predictions for flexible N- or C-terminal tails were removed from figures for clarity and space.

**Figure S1.**
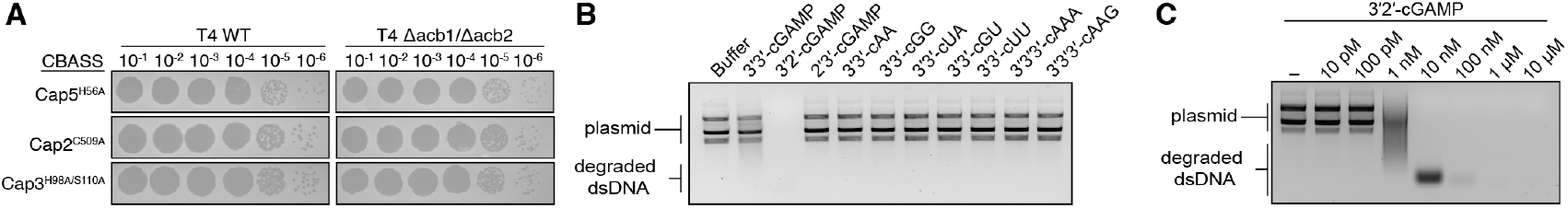
**(A)** Representative plaque assay depicting WT phage T4 or phage T4 Δacb1/Δacb2 plated on cells expressing CBASS operons with the indicated inactivating mutations. **(B)** Agarose gel depicting specific activation of Cap5 in the presence of 3′2′-cGAMP. All cyclic nucleotides were resuspended in water and used at a concentration of 1 μM. **(C)** Agarose gel depicting activation of Cap5 in response to titrating amounts of 3′2′-cGAMP.

**Figure S2.**
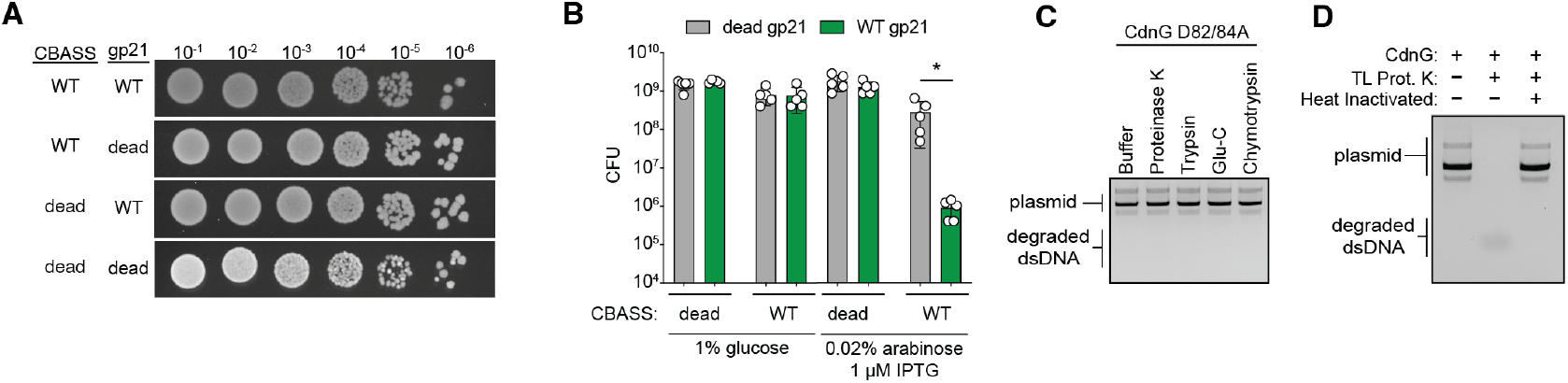
**(A)** Representative toxicity assay depicting serial dilutions of *E. coli* co-transformed with the indicated CBASS operon and phage T4 prohead protease gp21 construct and plated on LB supplemented with 1% glucose. For CBASS, ‘dead’ refers to *Ec*CdnG D82/84A mutations and for T4 gp21 ‘dead’ refers to H85A/S140A mutations. **(B)** Quantification of the data in Figure 2A and Figure S2A. **(C)** Representative agarose gel depicting no activation of Cap5 when *Ec*CdnG D82/84A is treated with the indicated proteases. **(D)** Representative agarose gel depicting activation of Cap5 and cleavage of indicator dsDNA when *Ec*CdnG is treated with thermolabile (TL) proteinase K but not when TL proteinase K has been heat-inactivated.

**Figure S3.**
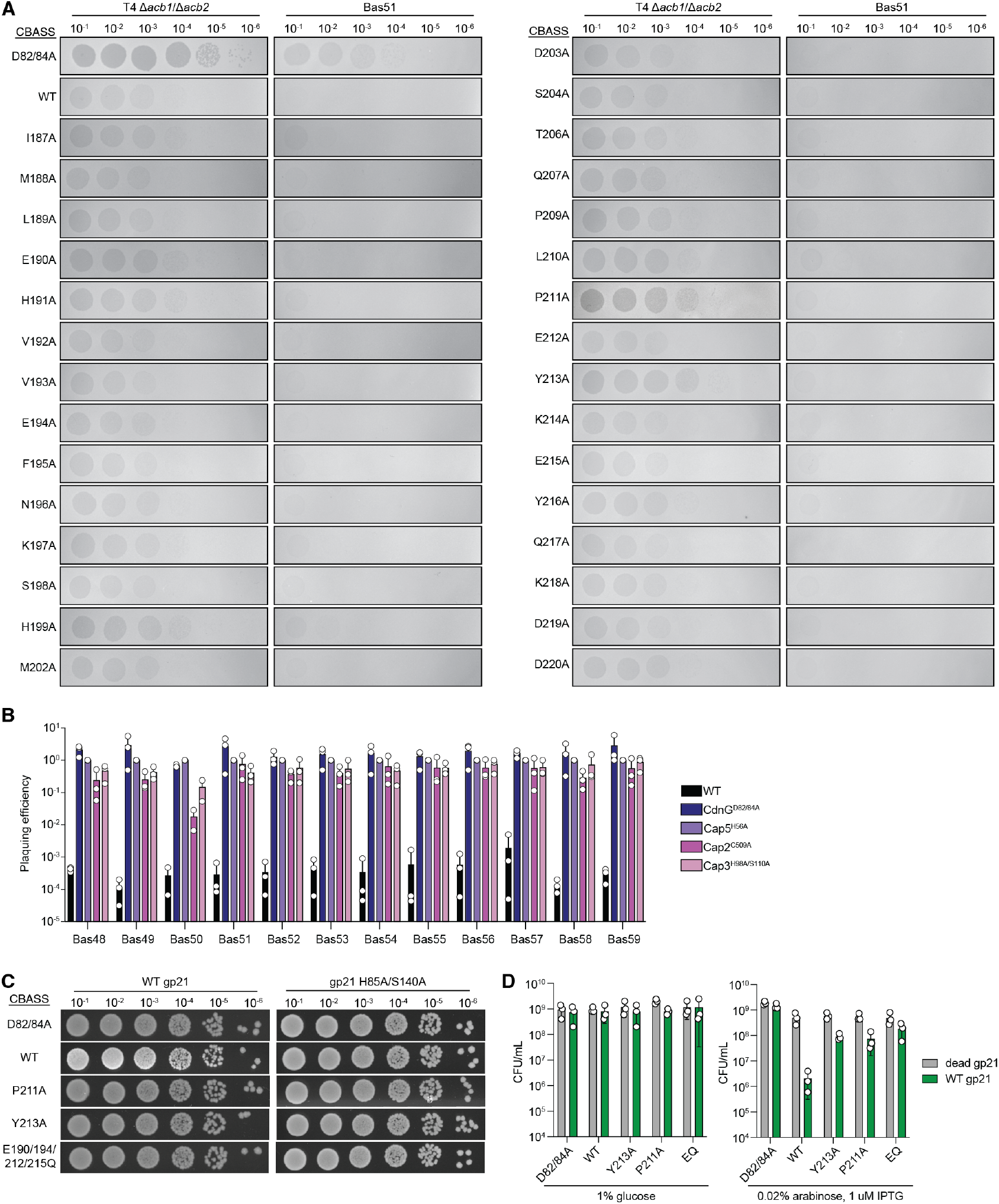
**(A)** Representative plaque assays depicting serial dilutions of phage T4 Δacb1/Δacb2 or phage Bas51 plated on cells expressing CBASS operons with the indicated *Ec*CdnG point mutations. **(B)** Quantification of Bas48–59 phage titers when plated on cells expressing the indicated CBASS operon. Data are normalized to titers when plated on the Cap5 H56A inactive operon. **(C)** Representative toxicity assay depicting serial dilutions of *E. coli* co-transformed with CBASS operons with the indicated *Ec*CdnG mutations and phage T4 prohead protease gp21 construct and plated on LB supplemented with 1% glucose. **(D)** Quantification of the data in Figure 3F and Figure S3C.

**Figure S4.**
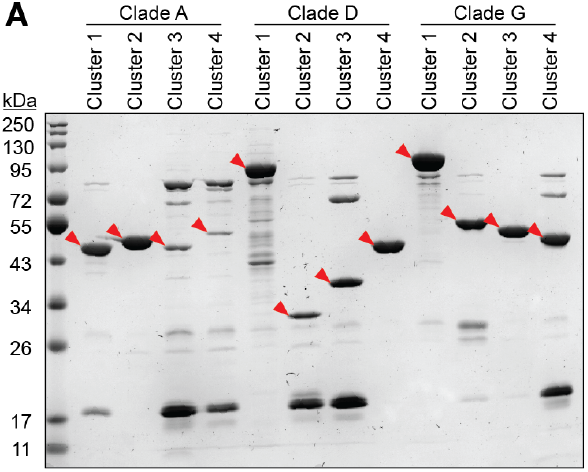
**(A)** SDS-PAGE analysis of final purified representative CD-NTase proteins from the indicated clusters used in Figure 4. Red arrows indicate bands that correspond to the predicted molecular weight of each CD-NTase. See Table S1.

**Table S1.** Details of CD-NTase enzymes used in Figure 4.

| Clade | Cluster number | NCBI Accession | Alternative names (if any) | Species | Whiteley et al CD-NTase number | Concentration used in Fig. 4 |
| --- | --- | --- | --- | --- | --- | --- |
| A | 1 | WP_065197317.1 |  | Stenotrophomonas maltophilia | 2135 | 50 nM |
| A | 2 | WP_001901330.1 | VcDncV | Vibrio cholerae | 2188 | 50 nM |
| A | 3 | WP_049588582.1 |  | Alteromonas macleodii | 2283 | 5 µM |
| A | 4 | WP_101217215.1 |  | Pseudoalteromonas | 2499 | 5 µM |
| D | 1 | WP_031517014.1 |  | Desulfotomaculum alkaliphilum | 4773 | 5 µM |
| D | 2 | WP_096173194.1 |  | Cohaesibacter sp. ES.047 | 4588 | 5 µM |
| D | 3 | SCJ44608.1 |  | uncultured Clostridium sp. | 4352 | 5 µM |
| D | 4 | WP_032676400.1 | EcCdnD02 | Enterobacter cloacae | 3920 | 50 nM |
| G | 1 | WP_085881083.1 |  | Roseisalinus antarcticus | 329 | 5 µM |
| G | 2 | NP_766712.1 | BdCdnG | Bradyrhizobium diazoefficiens USDA 110 | 884 | 50 nM |
| G | 3 | WP_000064266.1 | EcCdnG | Enterobacteriaceae | 71 | 50 nM |
| G | 4 | WP_093192452.1 |  | Thiocapsa sp. KS1 | 373 | 50 nM |

## Notes

### Competing Interest Statement

The authors have declared no competing interest.

### Funding

- NIH: 1DP2GM146250-01

